# The molecular architecture of engulfment during Bacillus *subtilis sporulation*

**DOI:** 10.1101/498220

**Authors:** Kanika Khanna, Javier Lopez-Garrido, Ziyi Zhao, Reika Watanabe, Yuan Yuan, Kit Pogliano, Elizabeth Villa

**Author notes:** Declaration of interests. To whom correspondence should be addressed: Elizabeth Villa, Tata Hall 5103, University of California San Diego, 9500 Gilman Drive, La Jolla CA 92093, Kit Pogliano, Natural Sciences Building 4109, University of California San Diego, 9500 Gilman Drive, La Jolla CA 92093.

## Abstract

The study of cell biology is limited by the difficulty in visualizing cellular structures at high spatial resolution within their native milieu. Here, we have visualized sporulation in *Bacillus subtilis* using cryo-electron tomography coupled with cryo-focused ion beam milling, a technique that allows the 3D reconstruction of cellular structures in near-native state at molecular resolution. During sporulation, an asymmetrically-positioned septum divides the cell into a larger mother cell and a smaller forespore. Subsequently, the mother cell phagocytoses the forespore in a process called engulfment, which entails a dramatic rearrangement of the peptidoglycan (PG) cell wall around the forespore. By imaging wild-type sporangia, engulfment mutants, and sporangia treated with PG synthesis inhibitors, we show that the initiation of engulfment does not entail the complete dissolution of the septal PG by the mother cell SpoIIDMP complex, as was previously thought. Instead, DMP is required to maintain a flexible septum that is uniformly and only slightly thinned at the onset of engulfment. Then, the mother cell membrane migrates around the forespore by forming tiny finger-like projections, the formation of which requires both SpoIIDMP and new PG synthesized ahead of the leading edge of the engulfing membrane. We propose a molecular model for engulfment membrane migration in which a limited number of SpoIIDMP complexes tether the membrane to and degrade the new PG ahead of the leading edge, thereby generating an irregular engulfing membrane front. Our data also reveal other structures that will provide a valuable resource for future mechanistic studies of endospore formation.

## Introduction

From an architectural point of view, bacterial cells are among the simplest forms of life on the planet. Their cytoplasm is typically devoid of membrane bound organelles, and their cellular morphology relies upon a semi-rigid peptidoglycan (PG) cell wall that imposes its shape on the malleable cell membrane(s). Bacterial cells are inflated by their high internal turgor pressure, which pushes the membranes against the cell wall, causing the PG to stretch and the cell to adopt its appropriate shape. Despite the apparent simplicity, studies in the past few decades have demonstrated that bacterial cellular architecture is far more complex than previously thought, in terms of both its ultrastructure and dynamic capabilities ^1–3^.

Endospore formation in *Bacillus subtilis* represents a striking example of the dynamic capabilities of bacterial cells, as it entails dramatic changes in cellular architecture. First, the division sites shift to polar positions, generating a sporangium comprised of two cells: a larger mother cell and a smaller forespore (Fig. 1A; ^4–6^). The polar septum traps the forespore chromosome, which is subsequently transported to the forespore by SpoIIIE, a membrane-anchored ATPase that assembles a translocation complex at septal midpoint ^7–10^. Chromosome translocation increases the turgor pressure in the forespore, causing it to inflate and expand into the mother cell ^11^. Simultaneously, the mother cell engulfs the forespore in a process that resembles eukaryotic phagocytosis (Fig. 1A). After engulfment, the forespore is fully enclosed within the cytoplasm of the mother cell, where it matures in a process that involves the synthesis of protective layers of cortex and coat, and the partial dehydration of the forespore cytoplasm. Finally, the mother cell lyses and the mature spore is released to the environment, where it remains dormant until the conditions are appropriate for germination.

**Figure 1.**
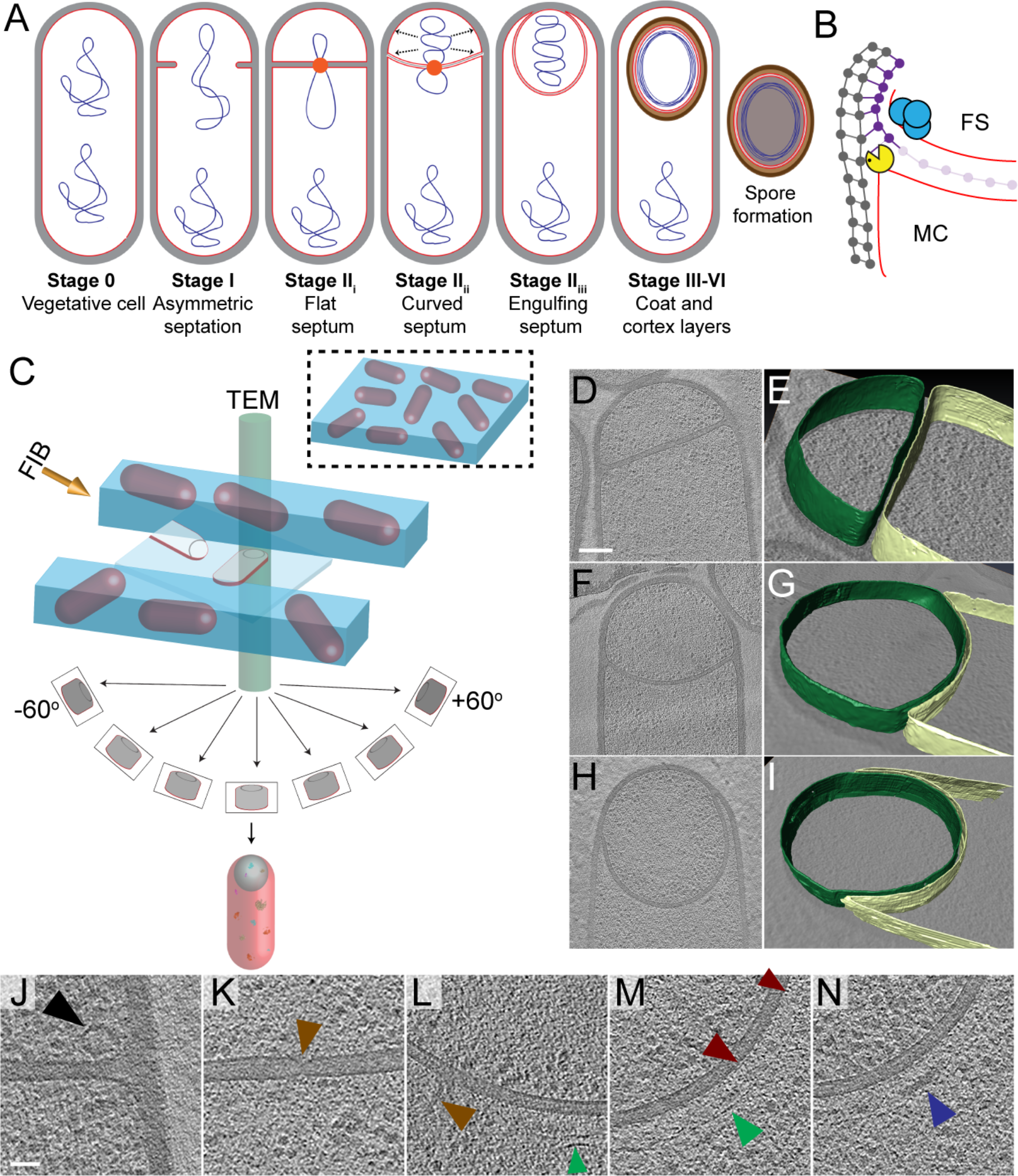
Visualizing the 3D architecture of engulfment during sporulation in *B. subtilis*. *(A)* Schematic illustrating the process of polar septation, chromosome translocation, engulfment and spore maturation. Membranes (red), PG (grey), chromosomes (blue), SpoIIIE (orange) and coat proteins (shades of brown) are highlighted. Outward arrows in the stage IIii forespore indicate increased turgor pressure due to chromosome translocation. *(B)* Revised engulfment model (cf. ref. 23): new PG (purple) is synthesized ahead of the leading edge of the engulfing membrane by forespore-associated PG biosynthetic enzymes (blue) and is subsequently degraded by DMP (yellow pacman), making room for the engulfing membrane to advance. The coordinated synthesis and degradation of PG at the leading edge of the engulfing membrane can move the junction between the septum (pink) and the lateral cell wall (grey) around the forespore. *(C)* Schematic illustrating cryo-FIB-ET methodology for bacterial samples. (See *Materials and Methods*). *(D-I)* Slices of tomograms representing different stages of engulfment: *(D)* polar septum (Stage IIi), *(F)* curved septum (Stage IIii) and *(H)* engulfing septum (Stage IIiii). Scale bar: 200 nm. The corresponding forespore (green) and mother cell (yellow) are annotated in *(E,G,I)* respectively. Scale bars have been omitted for these as perspective views of the cells are shown. *(J-N)* Zoomed-in slices of tomograms showing *(J)* a large ellipsoidal complex adjacent to the forespore membrane (black arrow), *(K)* putative SpoIIIE channel and *(L)* other channel (brown arrows), *(L,M)* coat filaments (green arrows), *(M)* basement coat layer (maroon arrow) and *(N)* amorphous coat proteins (purple arrow). Scale bar: 50 nm. See also Fig. S1.

Engulfment represents a major rearrangement of the sporangium, from two cells that lie side by side to a cell within a cell. Such rearrangement likely involves a profound remodeling of the PG cell wall around the forespore, which would otherwise constrain the movement of engulfing membrane. At the onset of engulfment, the engulfing mother cell membrane must circumvent the physical barrier posed by the septal PG in order to migrate around the forespore. This has led to the logical proposal that engulfment must entail the complete dissolution of the septal PG, a process often referred to as ‘septal thinning’ ^12–14^. This proposal was supported by early electron microscopy studies of fixed and stained sporangia showing that engulfment-defective mutants had thicker septa than wild type sporangia ^12,15,16^. Further studies showed that engulfment requires three mother cell proteins: SpoIID, SpoIIM and SpoIIP, which form a complex (DMP) with PG degradation activity ^14,17–19^. In principle, DMP could mediate the complete dissolution of the septal PG to remove the steric block to the movement of the mother cell membrane around the forespore.

The idea that septal PG is completely degraded has been more recently challenged by cryo-electron tomography (cryo-ET) images showing that a thin PG layer is present between the forespore and the mother cell membranes throughout engulfment ^20^. It has also been shown that DMP-mediated PG degradation is required and rate-limiting for membrane migration even after the septal barrier has been bypassed, suggesting that DMP plays a role separate from the dissolution of septal PG ^17,21^. In addition, the movement of the mother cell membrane also requires PG synthesis at the leading edge of the engulfing membrane ^22,23^. Based on these observations, we proposed a revised model for engulfment membrane migration in which coordinated PG synthesis and degradation at the leading edge of the engulfing mother cell membrane moves the junction between the septum and the lateral cell wall around the forespore, making room for the engulfing membrane to advance (^23^, Fig. 1B). This model eliminates the need for complete dissolution of septal PG and predicts that PG synthesis happens ahead of the leading edge of the engulfing membrane. Then, the mother cell DMP degrades this new PG to mediate membrane migration. However, due to the limited resolution of optical microscopy, conclusive evidence that PG synthesis occurs ahead of PG degradation is lacking.

Cryo-ET allows the visualization of three-dimensional (3D) architecture of bacterial membranes and cell wall in a hydrated near-native state that cannot be achieved by methods reliant on chemical fixation and staining ^24–26^. However, a limitation of cryo-ET is that the samples must be less than ~500 nm thick to obtain high-resolution tomograms, constraining its application to only a handful of bacteria that are either naturally slender, or, as in the case of *B. subtilis*, for which slender mutant strains are available ^20^. But the latter typically contain mutations in genes involved in PG metabolism and may not be ideal to study cell wall remodeling. Recent application of cryo-focused ion beam (cryo-FIB) milling has produced artifact-free thin sample sections of ~100-300 nm, which allows the acquisition of high-resolution tomograms of sections of wild type cells (Fig. 1C) ^27–29^. Cryo-FIB coupled with cryo-ET (cryo-FIB-ET) is therefore becoming the method of choice for studies of cellular architecture for both eukaryotic and prokaryotic cells ^11,30–32^.

Here, we have used cryo-FIB-ET to study sporulation in *B. subtilis*, revealing the different stages of engulfment with a resolution that has not been achieved previously. We have analyzed wild type sporangia, engulfment mutants, and sporangia treated with PG synthesis inhibitors to obtain new mechanistic insights into the PG transformations that occur during engulfment. First, we provide evidence that septal PG is not degraded completely at the onset of engulfment. Second, we show that during membrane migration, the newly synthesized PG deforms the forespore membrane ahead of the leading edge of the engulfing mother cell membrane, indicating that PG synthesis precedes PG degradation. Third, we observe that the mother cell membrane migrates around the forespore by forming tiny finger-like projections, the formation of which depends on both PG synthesis and degradation. The methodology, images and analysis presented here will provide valuable resources for future studies of spore assembly and other fundamental processes.

## Results

### Visualizing sporulation in wild type *B. subtilis* at molecular resolution

Recently, we used cryo-FIB-ET to illustrate the role of DNA translocation in inflating the forespore ^11^. These data confirmed the presence of a thin layer of PG between the forespore and the mother cell membranes in the wild type strain, as previously visualized by cryo-ET of a slender *ponA* mutant of *B. subtilis* ^20^. We expanded our cryo-FIB-ET studies to investigate the architecture of *B. subtilis* cells during different stages of sporulation (Fig. 1C; see *Materials and Methods*). We acquired high-quality data of wild type cells during engulfment (Stage II, Fig. 1D-I; Fig. S1A-L) and during later stages of sporulation, when the coat and the cortex were being assembled (Stages III-VI, Fig. S1T,U). Data of very late stages of sporulation (Fig. S1U) failed to provide high-resolution information inside the forespore, likely due to the dehydration of this cell, which increases sensitivity of cryo-ET samples to the electron beam ^33^.

Our data showed that the external cell wall of sporulating cells was ~20-30 nm thick, consistent with other EM studies ^20,24^. The polar septum formed close (within 500 nm) to one of the cell poles, and was initially flat (Fig. 1D,E, S1A-D). Eventually, the septum bent smoothly into the mother cell (Fig. 1F,G, S1E-H) and the mother cell membrane moved forward to engulf the forespore, at which stage the forespore was roughly rounded (Fig. 1H,I, S1I-L). Visual inspection of tomograms also revealed several structures that have not been characterized previously. Immediately after polar septation, we observed ellipsoidal complexes that were present only in the forespore, adjacent to the membrane and often close to the intersection between the septum and the lateral cell wall (Fig. 1J, S1M-O). The complexes were of roughly similar size, with a mean radius of ~45 nm (see *SI Materials and Methods*). The molecular identity of these structures remains undetermined. We also identified a region approximately in the center of a flat septum where the two membranes are closer together than elsewhere on the septum (14 nm *vs*. 23 nm, Fig. 1K, S1P,Q). This constriction may correspond to paired hexameric SpoIIIE channels ^10,34,35^. We also observed structures that appear to be channels crossing the septum (Fig. 1L) that may correspond to SpoIIQ-SpoIIIAH complexes ^36–38^. Next, we observed a basement coat layer adjacent to the outer forespore membrane that is likely comprised of SpoVM and/or SpoIVA (^39^, Fig. 1M). This layer is visible as an array of dots spaced ~4-6 nm apart, similar to that observed previously in *A. longum* sporulating cells ^40^. Moving outward, we observed a dense amorphous layer and a filamentous layer (Fig. 1L-N, S1R-S) that may contain CotE, SpoIVA and other coat proteins that are recruited to the outer forespore membrane during engulfment ^41^. Further studies are required to determine the molecular identity of these structures unambiguously.

### Septal PG is not completely degraded at the onset of engulfment

The cryo-FIB-ET images provided high-resolution details of the septum and the engulfing membrane. Hence, we focused on those details to obtain mechanistic insights about engulfment. The complete degradation of the septal PG during septal thinning has been traditionally considered a prerequisite for engulfment (13, 14, 17, 42). However, we observed a thin PG layer between the mother cell and the forespore membranes throughout engulfment in both wild type (Fig. 2A; Fig. S2A-C; ^11^) and a slender *ponA* mutant ^20^, suggesting that the septal PG is either not completely degraded or is quickly resynthesized after degradation. To probe this, we imaged engulfing sporangia after treatment with antibiotics that block PG synthesis: cephalexin and bacitracin (Fig. 2B,C, S2D-F). Blocking PG synthesis with either antibiotic blocks polar septation and engulfment membrane migration ^23^. We added the antibiotics two hours after inducing sporulation, when ~40-50% of the cells have undergone polar septation ^23^, and prepared samples for tomography either one or two hours later. If the septal PG was completely degraded, we would observe sporangia lacking PG between the mother cell and the forespore membranes after antibiotic treatment. However, if the septal PG was not completely degraded, the sporangia would show a layer of PG around the forespore, independent of antibiotic treatment. Indeed, we observed a thin PG layer in both untreated and antibiotic-treated sporangia (Fig. 2A-C, S2A-F), suggesting that the septum is not completely degraded at the onset of engulfment.

**Figure 2.**
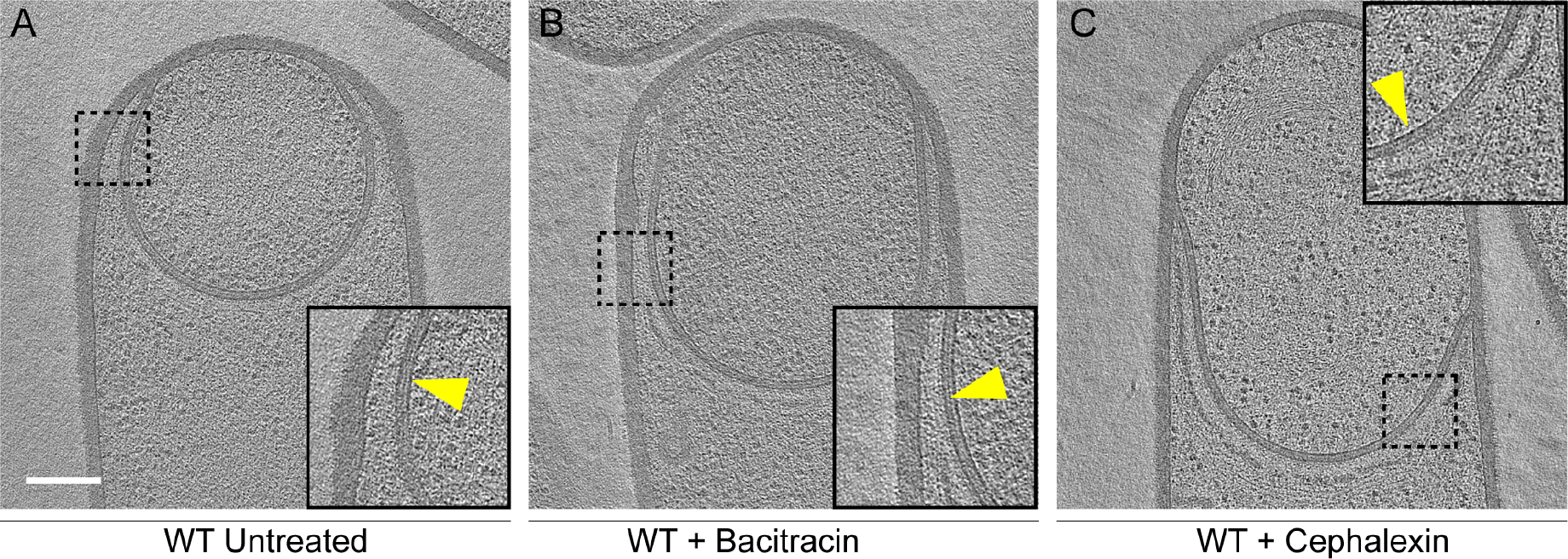
Septal PG is not completely degraded at the onset of engulfment. *(A-C)*Slices of tomograms of wild type sporangia that are *(A)* untreated, *(B)* bacitracin-treated and *(C)* cephalexin-treated during engulfment. The thin layer of septal PG is indicated by yellow arrows in zoom-in panels of the tomographic slices. Scale bar: 200 nm. See also Fig. S2.

### Septal thickness decreases slightly and uniformly across the entire septum during engulfment

The above observation prompted us to re-evaluate the process of septal thinning. The current model for septal thinning proposes that DMP initially localizes to the septal midpoint, where it starts degrading the septal PG as it moves towards the edge of the septal disk ^12,14,17,22^. This enzymatic septal thinning model predicts that, during the transition from flat to curved septa, the septum should be thinner in the middle than at the edges (Fig. 3A). To test this, we measured the distance between the forespore and the mother cell membranes across the length of the septum (*SI Materials and Methods*) for cells with flat (Stage II_i_), curved (Stage II_ii_) and engulfing (Stage II_iii_) septa (Fig. 3B-D, S3K-M). Sporangia with flat septa had an average septal thickness of ~23 nm (Fig. 3B,K) with 3 out of 5 septa being thicker at the middle (~28 nm) than at the edges (~22 nm) (Fig. 3B,J, S3K), contrary to what is proposed by the enzymatic septal thinning model. The septal thickness decreased by ~25% to ~18 nm during later stages of engulfment (Fig. 3C,D,K, S3L,M) and a thin layer of PG was observed in all septa (Fig. S2A-C). Importantly, septal thickness was uniform across the entire septum during later stages (Fig. 3C,D,J, S3L,M) and no septum was thinner in the middle than in the edges. These results show that the transition from thick to thin septum is homogenous, contrary to the prediction of the enzymatic septal thinning model.

**Figure 3.**
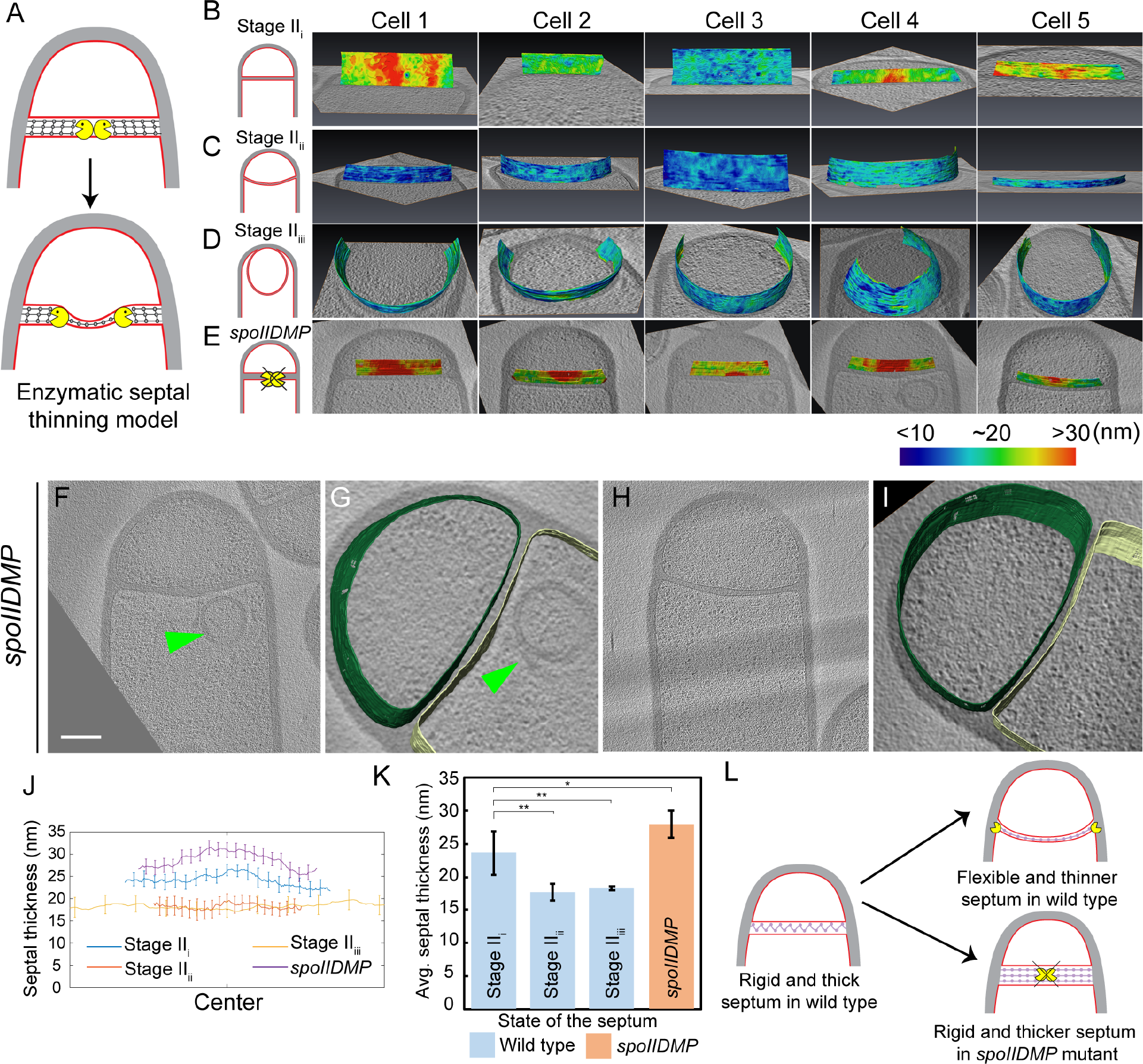
Septum is uniformly and only slightly thinned during engulfment and SpoIIDMP is required to maintain a thin flexible septum. *(A)* Schematic illustrating the model of septal thinning driven by septal PG (black meshwork) degradation by the DMP complex (yellow pacman) with membranes (red) and lateral PG (grey) highlighted. *(B-E)*Septal disc is color-coded according to the distance between the forespore and the mother cell membranes in five wild type sporangia at each stage: *(B)* flat, *(C)* curved and *(D)* engulfing septa, and *(E) spoIIDMP* mutant sporangia. Schematic representing the morphology of each cell type is shown on the far left. Scale bars have been omitted in distance plots as perspective views are shown. *(F,H)* Slices of tomograms of *spoIIDMP* mutant sporangia. *(G,I)* Annotated forespore (green) and mother cell (yellow) membranes for the tomograms in *(F,H)* respectively. Excess membrane accumulation is highlighted by green arrow in *(F,G)*. *(J)* Average thickness of the septum binned across the forespore surface for the cells shown in *(B-E)*. Error bars indicate standard deviation. For calculating average of *spoIIDMP* mutant sporangia, two additional tomograms shown in Fig. S3J were used. *(K)* Average septal thickness for wild type flat (Stage IIi), curved (Stage IIii) and engulfing (Stage IIiii) sporangia and *spoIIDMP* mutant sporangia. Error bars indicate standard deviation (n.s.: p > 0.05; *: p < 0.05; **: p < 0.01; ***: p < 0.001, unpaired t-test). *(L)* Schematic representing the role of DMP in septal thinning with membranes (red), lateral PG (grey), septal PG (pink), DMP complex (yellow pacman) highlighted. The rigid and thick septum in wild type can curve into the mother cell during engulfment but that of DMP mutant becomes even thicker and does not curve into the mother cell. See also Fig. S3 and S4.

### SpoIIDMP is essential to maintain a thin, flexible septum

Next, we tested whether DMP was required to mediate the slight thinning observed during the transition from flat to curved septum. To address this question, we measured septal thickness in DMP mutants. In single mutants lacking D, M or P, engulfment is blocked but the septum bulges towards the mother cell, which complicates the analysis (Fig. S3E-H, I,O,P). However, bulge formation is largely abolished in a triple mutant lacking functional versions of D, M and P simultaneously ^42^. Therefore, we imaged the DMP triple mutant by cryo-FIB-ET and measured its septal thickness (Fig. 3E-I, S3A-D,J,N).

Most sporulation septa of the DMP triple mutant were either flat or slightly curved into the mother cell (Fig. 3F-I, S3A-D). In some cells, we observed membrane accumulation in the mother cell (Fig. 3F,G, S3A,B) and small bulges approximately in the middle of the septum (Fig. S3C,D). Septal thickness ranged from ~25 nm to ~45 nm, with an average thickness of 28 nm (Fig. 3J,K, S3N), which is ~5 nm greater than that of wild type sporangia with flat septa (Fig. 3K). Importantly, there were no significant differences in septal thickness between flat and curved septa in DMP mutant sporangia (Fig. 3E, S3J,N), indicating that DMP is in fact necessary for the slight thinning of the septum observed in wild type sporangia.

Surprisingly, the thickness of individual DMP mutant septum was irregular across the septal length, with thicker regions of more than 45 nm, that were not observed in wild type cells (Fig. S3N). One possible explanation for this finding is that in the absence of DMP, proteins involved in PG synthesis persist at the septum, and their continued activity leads to thicker and less flexible septal regions (Fig. 3L). To test this model, we stained wild type and DMP mutant sporangia with BOCILLIN-FL, a fluorescent-derivative of penicillin V with affinity for multiple penicillin-binding proteins (PBPs) (^43^, *SI Materials and Methods*). We observed continuous fluorescent signal around the septa in both strains, but the signal was brightest at the leading edge in wild type sporangia (Fig. S4A, ^23^) and at different positions across the septum in DMP mutant sporangia (Fig. S4B). This mislocalization might allow ongoing synthesis of septal PG, leading to abnormally thick sporulation septa.

### PG is synthesized ahead of the leading edge of the engulfing membrane

Once the septum curves, the mother cell membrane starts to migrate around the forespore. Since the PG is not completely degraded, it will represent a major obstacle for the advancement of the engulfing membrane. To explain how cells overcome this hurdle, we previously proposed a conceptually new model for engulfment in which coordinated PG synthesis and degradation at the leading edge of the engulfing membrane moves the junction between the septum and the lateral cell wall around the forespore, making room for the engulfing mother cell membrane to advance (^23^, Fig. 1B). In this ‘make before break’ model, new PG would be synthesized ahead of the engulfing membrane by forespore-associated PG biosynthetic complexes and subsequently degraded by DMP. To test this model, we first focused on the shape of the forespore membrane opposing the leading edge of the engulfing mother cell membrane. In wild type sporangia, the forespore membrane was rounded immediately ahead of the engulfing membrane (Fig. 4A). This could be due to accumulation of additional PG at this site, which might push and deform the forespore membrane, introducing a broader curve. To confirm this, we analyzed the shape of the forespore membrane of sporangia in which PG synthesis was blocked with either bacitracin or cephalexin (*SI Materials and Methods*). The forespore membrane was less rounded and had a sharp corner (Fig. 4B, S5A) with a radius of curvature that was four times smaller than that of untreated cells (~27 nm vs. ~123 nm, Fig. 4C). These results indicate that new PG is indeed synthesized ahead of the leading edge of the engulfing membrane, and that it deforms the forespore membrane at the junction between the septum and the lateral cell wall.

**Figure 4.**
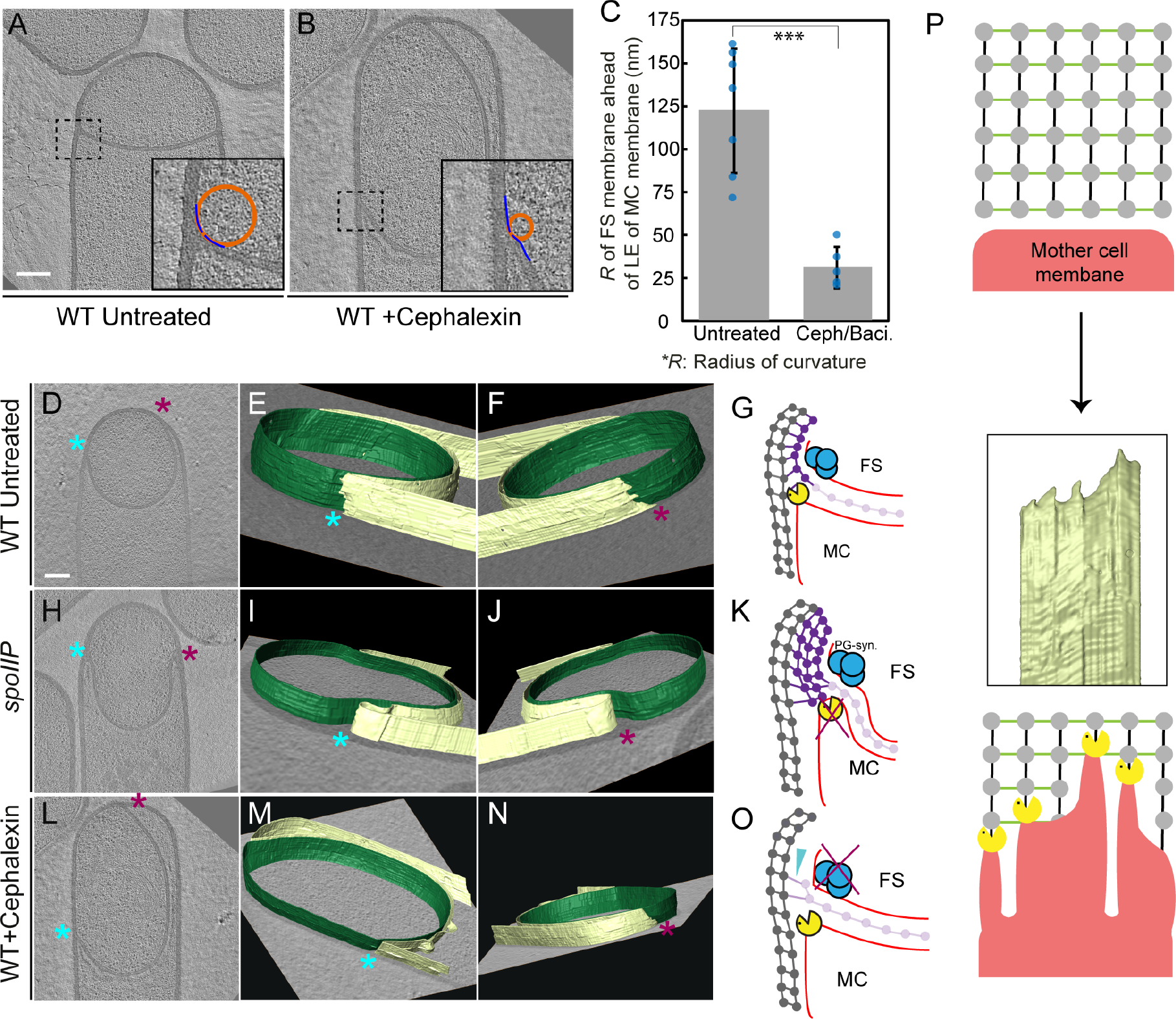
Architecture of the leading edge of the engulfing membrane. *(A,B)* Slices of tomograms of a wild type *(A)* untreated and *(B)* cephalexin-treated sporangia. The radius of curvature (orange circle) of the forespore membrane (blue spline) ahead of the leading edge of the engulfing membrane is highlighted in the respective insets. *(C)* Plot showing the average radius of curvature (nm) of the forespore membrane ahead of the engulfing membrane for untreated and antibiotic-treated sporangia. Blue dots indicate individual data points; black bars indicate the standard deviation (***: p< 0.001, unpaired t-test). *(D)* Slice of a tomogram of a wild type sporangium with annotated forespore (green) and mother cell (yellow) membranes shown in *(E, F)* as viewed from both left (blue asterisk) and right (maroon asterisk) sides. Similar labeling scheme is followed through *(H-N)*. *(G)* Schematic showing the localization of DMP PG degradation machinery (yellow pacman) and PG synthases (blue circles). Membranes (red), lateral PG (grey),septal PG (pink) and new PG (purple) are shown. *(H)* Slice of a tomogram of *spoIIP* mutant sporangium. *(I,J)* Annotated membranes for the tomogram shown in *(H)*. *(K)* Schematic representing a cell in which the DMP complex (yellow pacman) does not assemble. *(L)* Slice of a tomogram of cephalexin-treated sporangium. *(M,N)* Annotated membranes for the tomogram shown in *(L)*. *(O)* Schematic representing a cell in which PG synthesis (blue circles) has been inhibited. Scale bar for *(D,H,L)*: 200 nm. Scale bars have been omitted for surface rendered images owing to their perspective nature. *(P)* Model for mother cell membrane migration. DMP complexes (yellow pacman) present at different positions on the mother cell membrane (red) tether the membrane to the PG (grey) synthesized ahead. Due to a limited number of DMP complexes, the engulfing membrane may move forward in finger-like projections. This is indicated by annotated mother cell membrane (yellow) from Fig. 4F. See also Fig. S5.

### 3D architecture of the leading edge of the engulfing membrane

The new PG synthesized ahead of the leading edge of the engulfing membrane might interfere with the movement of the engulfing membrane, until it is degraded by DMP. To obtain insights about the movement of the engulfing membrane, we annotated the forespore and the mother cell membranes in our tomograms to visualize the architecture of the leading edge in 3D. Our data showed that the leading edge of the engulfing membrane in wild type sporulating cells moved around the forespore in tiny finger-like projections (Fig. 4D-G, S5B-D). The projections were ~10-30 nm wide and ~5-20 nm long, with significant variations from cell to cell. We hypothesized that those projections could be due to the uneven degradation of PG ahead of the leading edge of the engulfing membrane. To test this possibility, we imaged *spoIIP* mutant sporangia, which lack a functional DMP complex. As expected, the septum bulged towards the cytoplasm, but the mother cell membrane did not move forward in these cells. The 3D annotation of the membranes showed no membrane projections anywhere in the mother cell membrane (Fig. 4H-K, S5E-G). Also, no projections were observed in the DMP triple mutant, most of which did not form bulges (Fig. S5H-M). These findings suggest that SpoIIDMP is necessary for the formation of finger-like projections in the engulfing mother cell membrane.

We next tested whether PG synthesis was also required for the formation of finger-like projections. To study this possibility, we focused on sporangia in which PG synthesis was blocked using cephalexin. We have previously demonstrated that some cephalexin-treated sporangia show asymmetric engulfment, with the engulfing membrane moving around the forespore on one side but not on the other ^23^. Timelapse microscopy showed that this asymmetric engulfment is not due to differences in the speed of membrane migration on one side of the forespore compared to the other. Rather, the septal disk rotates, with one side of the disk extending and the other retracting ^23^. It is possible that in the presence of cephalexin, DMP cleaves the junction between the septum and the lateral cell wall, detaching the septal disk and allowing cup rotation. We used cryo-FIB-ET to compare the architecture of the leading side (the side of the membrane that moves around the forespore) and the lagging side (the side of the membrane that retracts) of the engulfing membrane (Fig. 4L-O). We observed finger-like projections in the leading side (Fig. 4N), but not in the lagging side (Fig. 4M).

Taken together, these results suggest that the finger-like projections at the leading edge of the engulfing membrane might be caused by tethering of the engulfing membrane to the newly synthesized PG via DMP (Fig. 4P).

## Discussion

In this study, we have visualized the developmental process of sporulation in *Bacillus subtilis* using state-of-the-art cryo-FIB-ET. We have obtained images of cells during different stages of sporulation at a resolution of a few nanometers, revealing new details about the architecture of spore assembly, as well as several hitherto unknown structures inside and around the developing spore (Fig. 1). Our results also provide mechanistic insights into engulfment, including the early step of septal thinning (Fig. 2, 3) and membrane migration (Fig. 4) which are captured in the model presented in Fig. 5.

**Figure 5.**
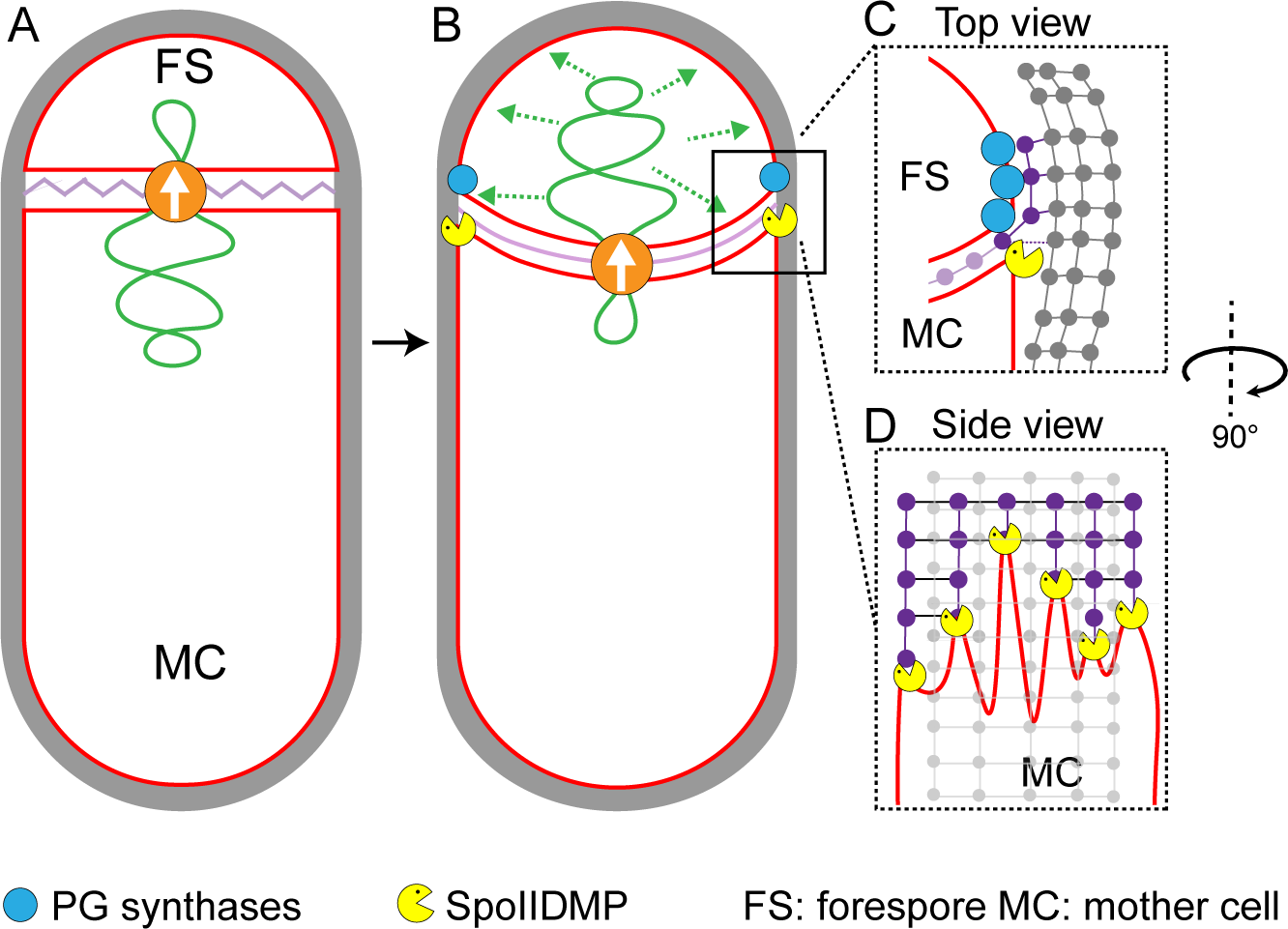
Model for septal thinning and membrane migration. *(A)* Schematic of a sporulating cell with a flat septum and relaxed septal PG (pink). Membranes (red), lateral PG (grey), SpoIIIE (orange) and forespore chromosome (green) are also highlighted. *(B)* As SpoIIIE translocated the chromosome to the forespore, the septal PG is stretched which may contribute to partial thinning of the septum. *(C)* Top view of the sporulating cell wherein coordinated PG degradation by DMP (yellow pacman) and PG synthesis by PG synthases (blue) at the leading edge provide room for mother cell membrane migration. *(D)* Side view showing the model for membrane migration with mother cell membrane (red), PG synthesized ahead of the membrane (purple) and lateral PG (grey) highlighted. During engulfment, DMP complexes present at different locations on the mother cell membrane may move forward by degrading the PG ahead and their rate-limiting activity may lead to the formation of tiny finger-like projections. FS: Foresepore; MC: mother cell.

We provide evidence that the septal PG is not completely degraded at the onset of engulfment. Instead, the septum gets slightly (~25%) thinner as it curves into the mother cell, with PG continuously present between the mother cell and the forespore membranes. In addition, the whole septum transitions from thick to thin homogenously, contrary to previous results that suggested that septal thinning started in the middle and progressed towards the edges. The homogenous thinning of the septum can be explained by our previous finding that septal PG is stretched as the forespore grows towards the mother cell due to increased turgor pressure caused by SpoIIIE-mediated chromosome translocation ^11^. Thus, septal thinning could simply represent a change in conformation of the septal PG, from a relaxed to a stretched state, triggered by the increased turgor pressure in the forespore. This is consistent with recent molecular dynamics simulations on gram-positive cell walls, which show that relaxed PG fragments are ~25% thicker than those in a strained conformation ^44^.

Our results also indicate that DMP is required to maintain a flexible septum that can curve as the forespore grows into the mother cell (Fig. 3). We found that the septa of DMP triple mutants have irregular thickness, and are on an average thicker than wild type septa (Fig. 3J,K). Since DMP is produced after polar septation, the septa of DMP mutant sporangia must have thickened after they have been formed. Thus, it is possible DMP prevents septal thickening by clearing PG synthases from the septum, where they would accumulate after polar septation. This model is consistent with the increased localization of PBPs throughout the septum in DMP mutants (Fig. S4). Nevertheless, we cannot rule out the possibility that DMP degrades the septal PG partially to generate a flexible septum. If this was the case, the partial degradation should happen simultaneously across the whole septum to enable the homogenous transition from thick to thin septum. However, since DMP is rate limiting for engulfment, it seems unlikely that there may be enough DMP complexes to mediate the homogeneous thinning of the septum.

Our data also provide insights into the mechanism of membrane migration during engulfment. Using fluorescence microscopy, we previously observed that new PG is synthesized at the leading edge of the engulfing membrane ^20,23^. We also showed that many forespore penicillin binding proteins (PBPs) can track the leading edge of the mother cell engulfing membrane ^23^, suggesting that PG synthesis at the leading edge of the engulfing membrane is carried out by forespore PBPs. The cryo-FIB-ET images presented here provide further support to this idea: comparing the architecture of the leading edge of the engulfing membrane between native sporangia and sporangia in which PG synthesis is blocked by antibiotics shows that new PG deforms and rounds the forespore membrane ahead of the leading edge of the engulfing membrane (Fig. 4A-C), indicating that new PG is synthesized by forespore PBPs immediately ahead of the leading edge of the engulfing cell membrane. We propose that DMP complexes at the leading edge of the engulfing membrane target this new PG for degradation, making room for the engulfing membrane to advance (Fig. 1B, 5C). This relatively simple model suggests that engulfment could have evolved by developing new mechanisms to spatially and temporally regulate the conserved protein machineries that synthesize and degrade peptidoglycan.

The 3D reconstruction of the leading edge of the engulfing mother cell membrane shows the presence of finger-like projections that resemble the filopodia of eukaryotic cells ^45^. In eukaryotic cells, these membrane projections are produced by cytoskeletal proteins, and the projections are typically a few micrometers wide. In contrast, no cytoskeletal elements contributing to engulfment have been described so far (or visualized in our tomograms) and the finger-like projections at the leading edge of the engulfing membrane are only a few nanometers wide. Instead, our results demonstrate that DMP is required for the formation of these finger-like projections. In addition, no finger-like projections are observed when the engulfing membrane retracts by inhibition of PG synthesis with cephalexin, suggesting that tethering the membrane to PG is required to maintain these projections. The most straightforward model to explain these results is that DMP tethers the engulfing membrane to the new PG ahead of the leading edge of the engulfing membrane, as it degrades the new PG to make room for the engulfing membrane to expand. DMP activity is rate limiting for membrane migration ^17,22^, suggesting that there is a discrete number of DMP complexes to remove the steric barrier posed by the newly synthesized PG ahead of the leading edge. The limited number of DMP complexes might cause the PG to be degraded irregularly, generating an uneven membrane front in the form of finger-like projections. The distance between the tips of the finger-like projections is ~5-20 nm, which would require that some DMP complexes are ahead of others by 2 to 8 glycan strands ^46^. From a functional perspective, we speculate that the finger-like projections could be compared to ‘friction ridges’, the minutely raised ridges of the epidermis that provide a grasping surface on our fingers ^47^. During engulfment, these membrane projections may provide the engulfing membrane a tighter lateral grip while moving around the forespore, hence serving as a ratchet that prevents backward movement of the membrane.

The application of cryo-FIB-ET has been instrumental in allowing us to visualize and accurately measure structural details of engulfing sporangia, which transpire at a scale of a few nanometers. Our studies support a mechanistic model for the enigmatic process of engulfment, while revealing new architectural details about engulfment and spore assembly, including intriguing new molecular structures that will require further study to unambiguously identify them. Our findings reveal details about sporulation at an unprecedented resolution and further illustrate the potential of cryo-FIB-ET to reveal critical new information about dynamic biological processes.

## Supporting information

## Acknowledgements

This work was supported by National Institutes of Health Director&’s New Innovator Award 1DP2GM123494 (E.V.), and the National Institutes of Health R01-GM057045 (E.V. and K.P.). We used the UC San Diego Cryo-EM Facility (supported by NIH grants to Dr. Timothy S. Baker and the Agouron Institute), and the San Diego Nanotechnology Infrastructure of UC San Diego (supported by the NSF grant ECCS-1542148). The authors would like to thank Antonio Martímez-Sánchez for useful inputs.

## Author contributions

K.K., J.L.G., K.P., E.V. designed research; K.K., J.L.G. performed research; K.K., J.L.G, Z.Z., Y.Y. analyzed data; R.W. assisted in cryo-EM; K.K., J.L.G. wrote original draft; all authors revised and edited the manuscript.

## Materials and Methods

### Strains and culture conditions

We used *Bacillus subtilis* PY79 for all data acquisition. The strains were grown in LB plates at 30°C. The bacteria were first grown in ¼ diluted LB to OD_600_ ~0.5-0.7. Sporulation was then induced by resuspension in A+B media at 37°C. For tomography, we collected wild type sporulating cells 1.5-3 hours after sporulation induction. For cells treated with antibiotics, 50 μg/ml of bacitracin or 50 μg/ml of cephalexin was added two hours after induction of sporulation and samples for tomography were collected either one or two hours later respectively. For *spoIIP* and *spoIIDMP* mutant sporangia (*SI Materials and Methods*), cells were collected 2.5 hours after induction of sporulation for tomography.

### Cryo-FIB-ET and Image Processing

Vitrified bacterial samples forming a monolayer on the grids were mounted into modified autogrids (Max Planck Institute for Biochemistry) for milling inside a Thermo Scientific™ Scios™ DualBeam™ (cryo-FIB/SEM) (*SI Materials and Methods*). 100-250 nm thin sections or lamellae (12-15 μm in width) were then prepared using rectangular milling patterns and beam current of 0.1 nA or 0.3 nA for rough milling and 10 pA or 30 pA for fine milling as described in ^30^. Tilt series were collected in a 300-keV Tecnai G2 Polara (Thermo Scientific&™) equipped with a K2 Summit direct detection camera (Gatan) and a post-column Quantum energy filter (Gatan). The samples were usually tilted from −66° to &#x002B;66° (the range per tilt series depending on the quality of the lamellae) with an increment of 1.5° or 2°. The tilt series were acquired using SerialEM ^48^. The images were recorded at a defocus of −5 to −6 μm at nominal magnifications of 34,000 (pixel size: 0.61 nm) or 41,000 (pixel size: 0.49 nm) with a cumulative dose of ~ 60-130 e^−^/Å^2^. The patch-tracking feature of IMOD was used to reconstruct the tilt-series ^49^. TomosegmemTV ^50^ was used for semi-automatic segmentation of membranes followed by manual refinement in the Amira™ software package.

