## Supplementary material for "The molecular architecture of engulfment during Bacillus *subtilis sporulation*"

**This PDF file includes:**

Supplementary text
Figs. S1 to S5
Captions for movies S1 to S6
References for SI reference citations

**Other supplementary materials for this manuscript include the following:**

Movies S1 to S6

**Supplementary Information Text**

**Materials**

| Strain | Genotype or description | Reference, source or construction |
|---|---|---|
| PY79 | Wild type | (1) |
| KP513 | *ΔspoIIP::tet* | (2) |
| KP4188 | *spoIID298*, *spoIIM-mls*, *ΔspoIIP::tet* | (3) |

**Methods**

**Additional details of sample preparation for cryo-FIB-ET**

7 µl of diluted liquid culture was deposited onto glow-discharged (using Pelco easyGlow™) holey carbon coated QUANTIFOIL R 2/1 200 mesh copper grids. Manual blotting was performed using Whatman No. 1 filter paper from the reverse side to remove excess resuspension media such that cells form a monolayer on the grids. They were then immediately plunge-frozen into a liquid ethane/propane mixture cooled by liquid nitrogen using a custom-built vitrification device (Max Planck Institute for Biochemistry, Munich). These grids were then stored in storage boxes in liquid nitrogen until further use.

Images for later stages of sporulation (Fig. S1U) were acquired at nominal magnification of 22,500 (pixel size: 0.92 nm) as these samples were highly sensitive to radiation damage.

**Measuring mean radius of ellipsoidal complexes**

The mean radius (r) of an ellipse is given by:

$$r = \sqrt{\frac{a^2 + b^2}{2}}$$

where *a* is the semi-major axis of the ellipse and *b* is the semi-minor axis of the ellipse. For ellipsoidal complexes observed in our tomograms (Fig. 1I, S1M-O), average value of *a* is ~45 nm and that of *b* is ~42 nm.

**Calculating septal distances**

'Surface distance' feature of Amira$^{TM}$ was used to calculate the septal distances between the forespore and mother cell membranes. The septal-disc was color coded according to these values (Fig. 3B-E, S3I,J). 5 tomograms each of wild type cells representing different stages of engulfment (flat, curved and engulfing septa) were used for analysis. For engulfment mutants, 5 *spoIIP* sporangia and 7 *spoIIDMP* sporangia were analyzed. To get the linear profiles of the distances as shown in Fig. S3K-O, the data was grouped into smaller bins of approximately equal area for each tomogram. Then the average of the distance within a bin was used to represent the distance between the membranes at that location of the bin. Fig. 3J represents an average of all the profiles for each cell type in Fig. S3K-O.

**Fluorescence microscopy**

Cells were visualized on an Applied Precision DV Elite optical sectioning microscope equipped with a Photometrics CoolSNAP-HQ2 camera and deconvolved using SoftWoRx v5.5.1 (Applied Precision). For experiment outlined in Fig. S4, the median focal plane of the image is shown. Membranes were stained with 0.5 μg/ml of FM4-64 (Thermo Fisher Scientific) that was added directly to 1.2% agarose pads prepared using sporulation resuspension medium. 10 μg/μl of BOCILLIN-FL was added to 1 ml of culture aliquoted at ~2.5 hours after sporulation induction (at 37$^{o}$C) and washed with sporulation resuspension medium three times. 12 μl of washed cells were then transferred to agarose pads for imaging.

**Calculating radius of curvature**

To calculate radius of curvature (Fig. 4A-C, S5A), a slice was taken approximately from the center of z-stack for each of the tomograms. 'Measure spline curvature' feature of sabl_mpl (4) was then used to plot radii of curvatures around the forespore membranes just ahead of the leading edge for 7 native sporangia and 5 antibiotic-treated sporangia wherein membrane migration appears to be blocked.

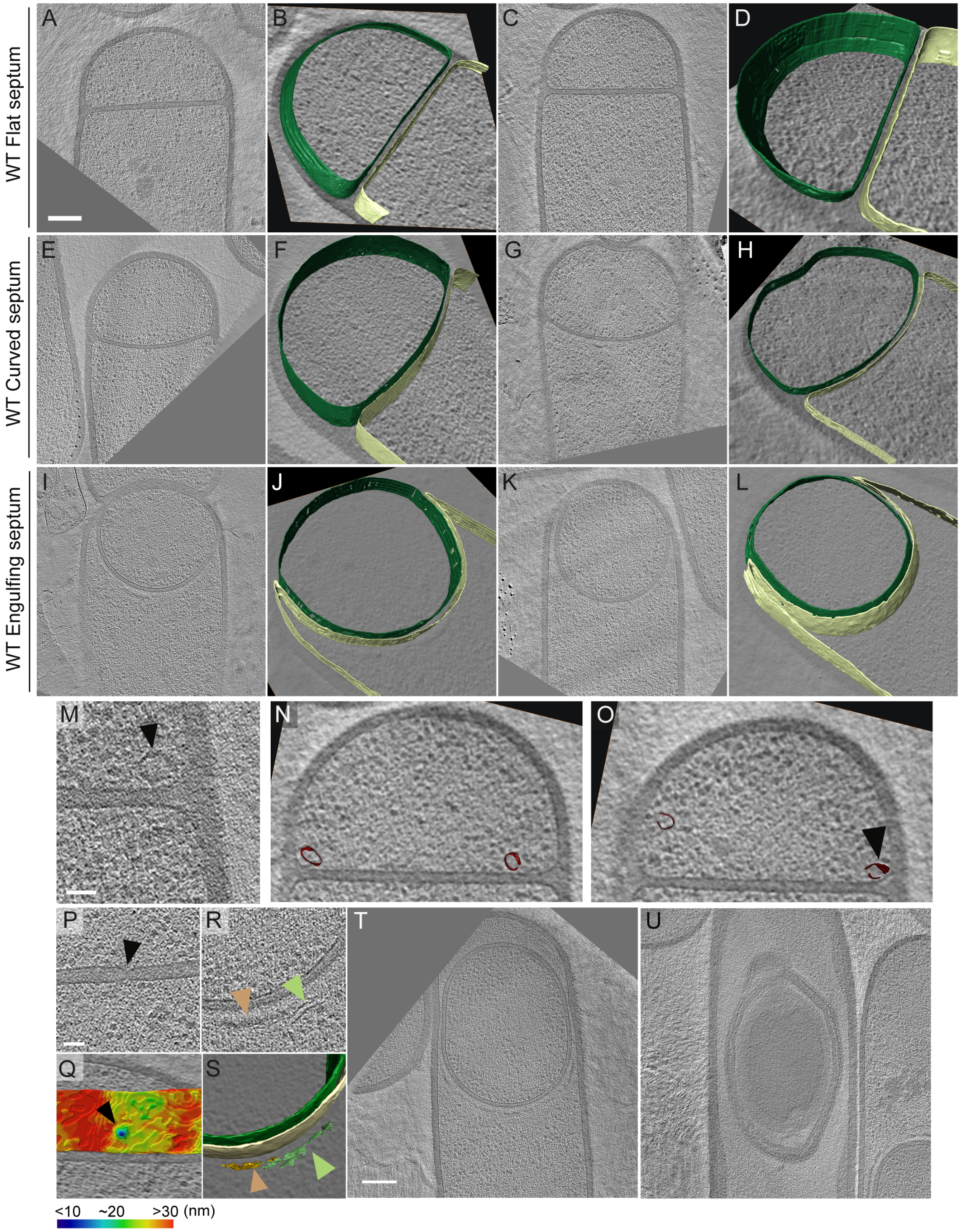


**Figure S1.** *(A-L)* Additional slices of cryo-electron tomograms representing different stages of engulfment in the wild type. The forespore (green) and the mother cell (yellow) membranes are annotated and shown adjacent to each tomogram. *(A-D)* represent flat septum (Stage $II_i$) sporangia, *(E-H)* represent curved septum (Stage $II_{ii}$) sporangia and *(I-L)* represent engulfing septum (Stage $II_{iii}$) sporangia. Scale bar: 200 nm for all the tomogram slices from *(A)* through *(L)*. *(M)* Ellipsoidal structure seen in the forespore during early stages of septum formation (black arrow) as shown in Fig. 1I. Scale bar: 50 nm. *(N,O)* 2 different slices of the same tomogram showing the 3D annotation of the ellipsoidal structures (maroon) seen in the tomogram in *(M)*. The structure highlighted in *(M)* in black arrow is pointed out at in *(O)*. *(P)* Slice of a tomogram with putative SpoIIIE channel highlighted (black arrow) as in Fig. 1J and the corresponding color-coded distance of the forespore membrane from the mother cell membrane is shown in *(Q)*. The putative channel corresponds to a septal thickness of ~14 nm. *(R)* Slice of a tomogram with coat filaments (green arrow) and amorphous dense coat layer (brown arrow) highlighted. The corresponding 3D annotations are shown in *(S)*, along with the forespore membrane (green) and the mother cell membrane (yellow). Scale bar for *(P,R)*: 50 nm. *(T,U)* Slices of tomograms representing later stages of sporulation when the forespore is completely inside the mother cell. *(T)* represents an earlier stage than *(U)*. A dense spore with coat and cortex layers is observed in *(U)*. Scale bar for *(T,U)*: 200 nm. Scale bars have been omitted for all segmented images as cells are being viewed in perspective mode.

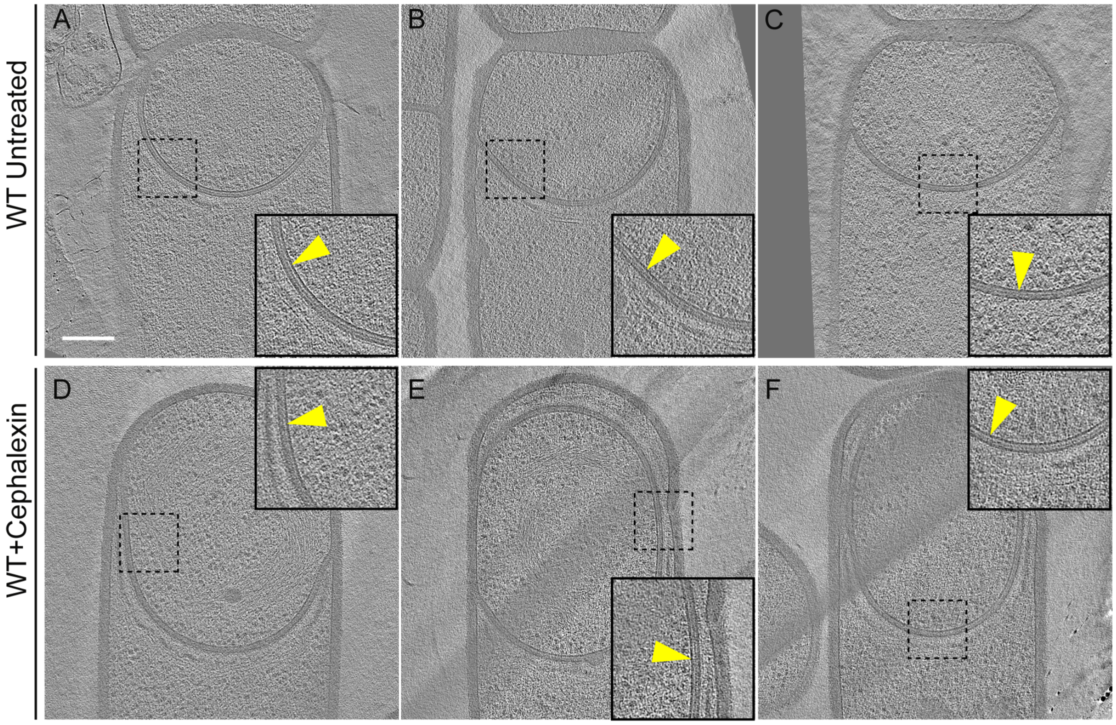


**Figure S2.** *(A-C)* Additional slices of tomograms of wild type untreated sporangia, and of *(D-F)* cephalexin-treated wild type sporangia. The thin layer of PG between the forespore and the mother cell membranes is indicated by yellow arrows in the zoom-in panels for each tomographic slice. Scale bar: 200 nm.

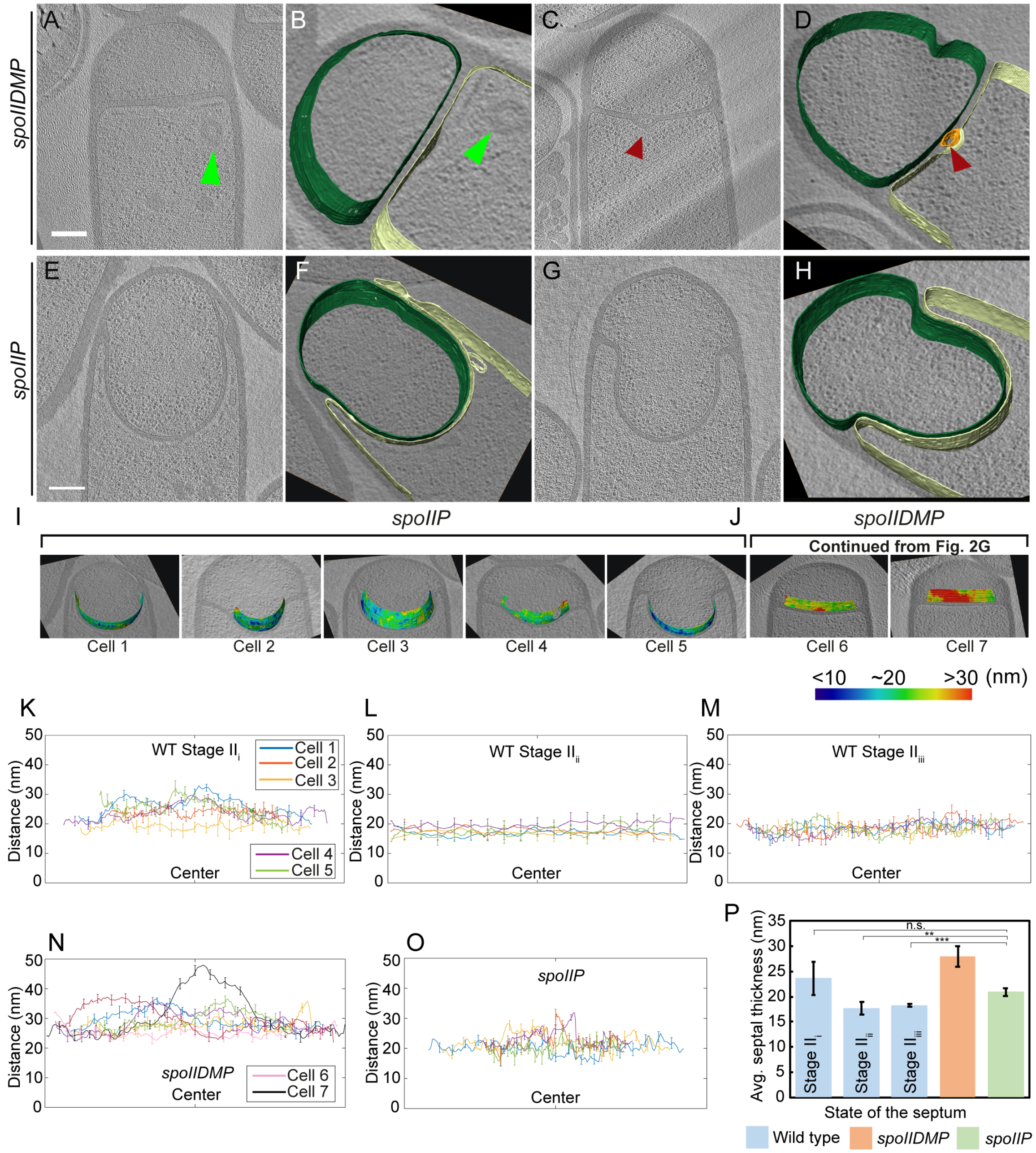


**Figure S3**. *(A-D)* Slices of tomograms of *spoIIDMP* sporangia with annotations of the forespore membrane (green) and the mother cell membrane (yellow) adjacent to each tomogram. Bulges are indicated by green arrows in *(A,B)* and red arrows in the middle of the septum in *(C,D)*. *(E-H)* Slices of tomograms of *spoIIP* sporangia with annotations of the forespore membrane (green) and the mother cell membrane (yellow) adjacent to each tomogram. Scale bars: 200 nm. *(I,J)* Color-coded distance of the forespore and the mother cell membranes along the length of the septum for *(I) spoIIP* sporangia and *(J)* the remaining two *spoIIDMP* sporangia not included in Fig. 3E. Scale bars are omitted for all segmented images due to their perspective nature. *(K)* Average thickness of the septum binned across the forespore surface for five WT Stage $II_{i}$ sporangia shown in Fig. 3B. *(L)* Average thickness of the septum binned across the forespore surface for the five WT Stage $II_{ii}$ sporangia shown in Fig. 3C, *(M)* Average thickness of the septum binned across the forespore surface for the five WT Stage $II_{iii}$ sporangia shown in Fig. 3D. *(N)* Average thickness of the septum binned across the forespore surface for seven *spoIIDMP* sporangia shown in Fig. 3E and Fig. S3J. *(O)* Average thickness of the septum binned across the forespore surface for five *spoIIP* sporangia shown in Fig. S3I. *(P)* Average septal thickness for wild type stages $II_{i\text{-}iii}$, *spoIIDMP* and *spoIIP* sporangia. Error bars indicate standard deviation (n.s.: $p > 0.05$; *: $p \leq 0.05$; **: $p \leq 0.01$; ***: $p \leq 0.001$, unpaired t-test).

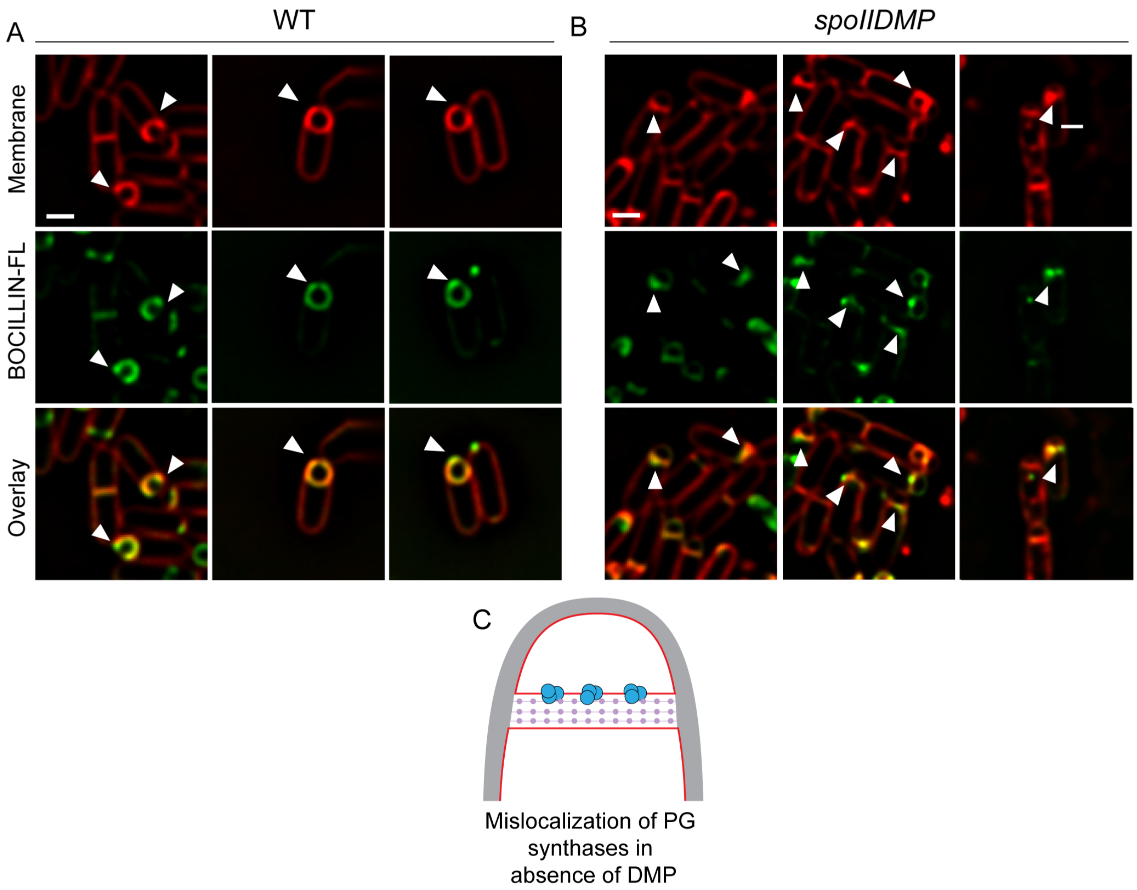


**Figure S4**. *(A)* Wild type sporulating cells stained with a green fluorescent derivative of penicillin V (BOCILLIN-FL). Membranes were stained with FM 4–64 (red). White arrows point at cells with enhanced fluorescent signal at the leading edge of the engulfing membrane. *(B) spoIIDMP* sporangia stained with BOCILLIN-FL (green). Membranes were stained with FM 4–64 (red). White arrows point at cells with enhanced fluorescent signal in regions of the septum. Scale bars: 1 μm. *(C)* Schematic to explain accumulation of PG synthases in the septum (blue circles) in absence of DMP as opposed to being localized at the leading edges as in wild type sporangia. Membranes (red), lateral PG (grey) and septal PG (pink circles) are also highlighted.

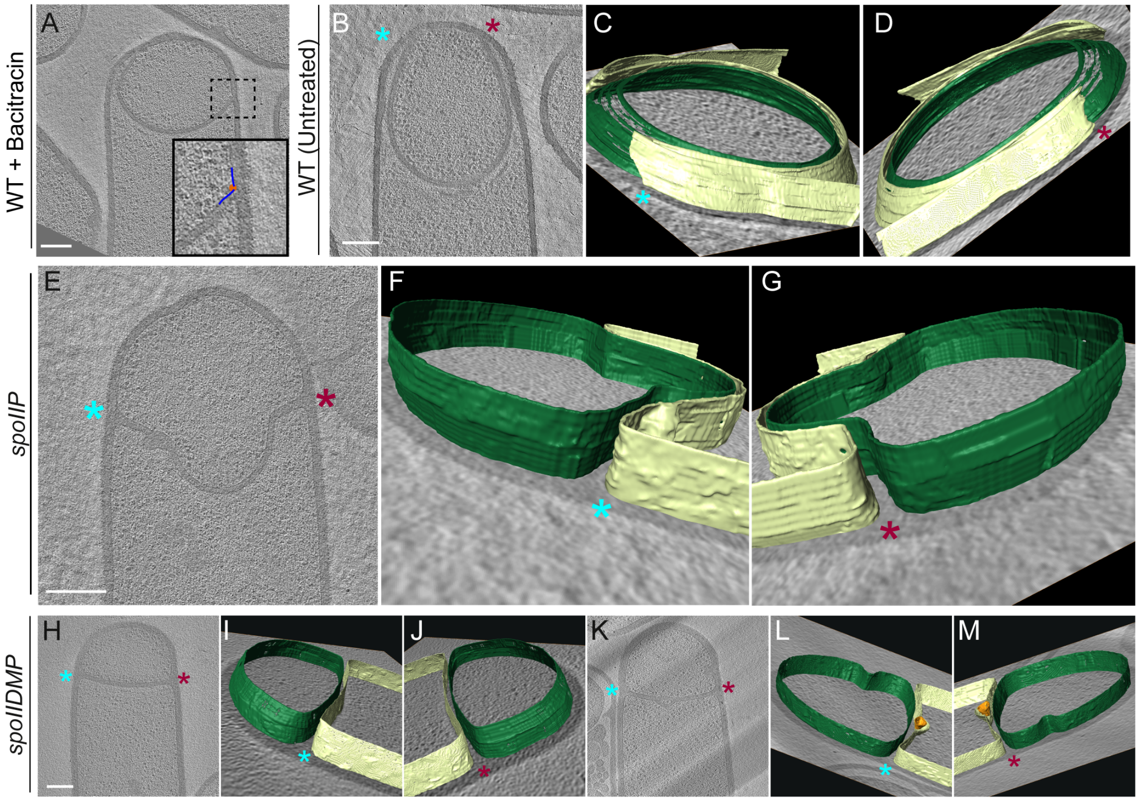


**Figure S5.** *(A)* Slice of a tomogram of a WT sporangium treated with bacitracin with the radius of curvature of the forespore membrane ahead of the leading edge of the engulfing membrane highlighted. *(B)* Slice of an additional tomogram of a wild type sporangium. *(C,D)* Annotation of membranes for the tomogram shown in *(B)*. Forespore membrane is shown in green and mother cell membrane in yellow and both sides of the membrane are indicated by blue and maroon asterisks in *(C,D,F,G,I,J,L,M)* and in their corresponding tomogram slices in *(B,E,H,K)* respectively. *(E)* Slice of an additional tomogram of *spoIIP* sporangium. *(F,G)* Annotation of membranes for the tomogram shown in *(E)*. *(H,K)* Slices of tomograms of *spoIIDMP* sporangia. *(I,J)* Annotation of membranes for the tomogram shown in *(H)*. *(L,M)* Annotation of membranes for the tomogram shown in (K). A bulge in the middle of the septum in *(K)* is highlighted in orange in *(L)* and *(M)*. Scale bars for all tomographic slices: 200 nm. Scale bars are omitted for annotated panels that are shown in perspective mode.

**Movie S1.** Movie showing slices of cryo-electron tomogram of *B. subtilis* WT sporulating cell (flat septum, Stage $II_i$) shown in Fig. 1D.

**Movie S2.** Movie showing slices of cryo-electron tomogram of B. subtilis WT sporulating cell (curved septum, Stage $II_{ii}$) shown in Fig. 1F.

**Movie S3.** Movie showing slices of cryo-electron tomogram of B. subtilis WT sporulating cell (engulfing septum, Stage $II_{iii}$) shown in Fig. 1H.

**Movie S4.** Movie showing slices of cryo-electron tomogram of SpoIIDMP mutant cell shown in Fig. S2C.

**Movie S5.** Movie showing slices of cryo-electron tomogram of SpoIIP mutant sporangia shown in Fig. 4H.

**Movie S6.** Movie showing slices of cryo-electron tomogram of wildtype sporangia treated with cephalexin shown in Fig. 4B.
